# Selective engagement of the primate orbitofrontal cortex during value-based but not perceptual decisions

**DOI:** 10.1101/2025.11.17.688922

**Authors:** Morgan G. Moll, Ryan K. Read, Joni D. Wallis, Thomas W. Elston

**Affiliations:** Department of Neuroscience, University of Texas at Austin, Austin, TX 78712; Department of Neuroscience, University of California at Berkeley, Berkeley, CA 94720

## Abstract

A fundamental question in neuroscience is whether the brain uses specialized sub-systems for different types of decisions or relies on a unified decision-making network. The orbitofrontal cortex (OFC) provides an ideal test case for this question: it has a well-established role in value-based decisions but it remains unknown whether this reflects functional specialization or participation in a broader, general decision network. To distinguish between these possibilities, we used Neuropixels to monitor large ensembles of OFC neurons as a monkey performed both value-based and perceptual decision tasks. Consistent with prior reports, OFC was robustly engaged during value-based decisions. In contrast, OFC was minimally engaged during perceptual decisions, with no significant encoding of task parameters at either the single neuron or population level. This highlights the functional specialization of the OFC for value-based decisions and suggests that the different cognitive demands underlying different types of decisions recruit distinct neural circuits.

## Introduction

The orbitofrontal cortex (OFC) plays a critical and causal role in value-based decision making. Damaging or inactivating OFC severely impairs value-based choices in both humans and animals (1–8). Neurophysiological studies revealed that OFC neurons encode the values of choice options during decision-making (9–11). Recent work suggests that OFC’s role in value encoding may reflect a more general function of simulating and comparing potential choice outcomes (12–14). In this framework, value is related to predicting and comparing the expected outcomes of different decisions. While OFC’s role in value-based choices is well-established, whether it plays a more general role in decision-making remains unclear.

Different types of decisions place fundamentally different demands on cognitive processing. Value-based decisions, such as choosing between different food options or investment strategies, require the brain to compute and compare the subjective worth of alternatives. In contrast, perceptual decisions, such as categorizing the direction of motion or identifying an object, involve extracting and reporting information from sensory input without necessarily weighing subjective preferences. Interestingly, human patients with selective ventromedial frontal damage show impaired preference judgments while their perceptual judgments remain intact (15). In contrast, patients with lateral prefrontal damage (as well as healthy controls with no brain damage) were unimpaired in both preference and perceptual judgements. This suggests that OFC may be selectively critical for value-based but not perceptual decisions and, more broadly, raises the hypothesis that different neural circuits are specialized for different types of decisions.

To test this hypothesis, we recorded neural activity from the OFC of a non-human primate trained to perform both value-based and perceptual decision tasks. If the brain utilizes a general, unified decision network, then OFC – given its established role in value-based decisions – should be engaged across both value-based and perceptual decisions. Conversely, if specialized decision systems are recruited according to cognitive demands, then OFC should be engaged during value-based but not during perceptual decisions.

## Methods and Materials

### Experimental subject details

All procedures were carried out as specified in the National Research council guidelines and approved by the Animal Care and Use Committee at the University of California, Berkeley. Both studies were performed on the same male rhesus macaque (subject K) aged 5 years and weighing 12 kg at the time of recordings. During the experiment, the subject sat head-fixed in a primate chair (Crist Instruments, Hagerstown, MD) and eye movements were tracked with an infrared system (SR Research, Ottawa, Ontario, CN). Stimulus presentation and behavioral conditions were controlled using the NIMH MonkeyLogic Matlab toolbox. The subject was implanted with a unilateral recording chamber that allowed access to the OFC.

### State-dependent valuation task

Monkeys were trained to perform a state-dependent value-based choice task involving 8 reward-predictive pictures (9,16). The value-mappings associated with each image depending on the cued task state. Stimuli were presented on a monitor positioned approximately 30 cm from the monkey. Subjects self-initiated trials by fixating on a white dot for 500 ms. Following fixation, a state cue indicated whether the upcoming choice should be evaluated according to the value scheme associated with state A, B, or C. Two distinct cues were used for each state to dissociate neuronal responses to visual properties from cue meaning. After 500 ms, the state cue disappeared, and the subject maintained central fixation for a further 100 ms. Then, the choice options were presented. On 75% of trials two options were shown (free choice trials), while on 25% of the trials there was only one option (forced choice trials). The presence of the forced choice trials was to ensure that all reward options were regularly experienced. Subjects reported their decisions by shifting their gaze to the chosen target location and fixating for 300 ms. Reward values corresponded to different amounts of apple juice delivered via a peristaltic pump.

For states A and B, the same choice stimuli were used, but the associated rewards differed depending on the cued state. For example, during a trial cued for State A, the “postbox” target would yield 1 drop of juice, while if state B were cued, it would instead be worth 4 drops of juice. State cues were pseudorandomized across trials. State C used a different set of choice options and had its own unique value schema that was not in conflict with the others. This control condition allowed for us to observe the decision processing without a conflicting value scheme.

### Perceptual categorization task

The subject was also trained on a perceptual decision making task which involved categorizing either the speed or direction of motion in a random-dot kinematogram (hence “motion stimulus”). Analogous to the state-dependent valuation task, subjects were cued whether to categorize according to speed or motion on each trial. Subjects initiated trials by fixating on a small white dot for 500 ms. They were then shown a cue for 600 ms indicating whether to attend to the speed or direction of the upcoming motion stimulus.

Like the state-dependent valuation task, two cues were used to indicate whether to direct attention to speed or direction in order to dissociate neural task representations from the visual features of the cues. After the cue and a 500 ms delay, the subject viewed a large motion stimulus (10 degrees of visual angle in radius) that varied in both speed (fast vs. slow) and direction (up vs. down). We associated each category with a different reward amount such that correctly categorizing either fast motion or downward motion led to a large juice reward (5 drops of apple juice) whereas correctly categorizing either slow motion or downward motion led to a smaller reward (3 drops of apple juice). Incorrect categorization led to no reward. We did this to test whether OFC encoded the expected reward associated with a given perceptual category.

Subjects reported their decisions during a dedicated choice epoch, which followed a 500 ms delay after the motion stimulus. The monkeys were trained to report fast motion and downward motion by saccading to a small speckled circle stimulus and, conversely, were trained to report slow motion or upward motion by saccading to a pink circle stimulus. Both speed and direction were present in every trial and we over-represented the “conflict” trials where the animal categorized the same motion stimulus differently depending on whether it was currently categorizing speed or direction. These “conflict” trials accounted for 70% of trials in a single behavioral session.

### Behavioral Analysis

We sought to compare the behavioral performance of the subject for both tasks. We began by looking at the subject’s ability to choose the correct option for different value combinations across each state. For the state-dependent value task, we calculated the probability of choosing the left option as a function of the value difference between the left and right options for each state. We then used a logistic regression model where the dependent variable was a binary indicator of left/right choice and the independent variables were task state (1,2, 3) and value difference (left minus right; +/- 1-3) (LR 1).

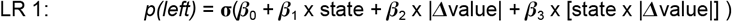

For the perceptual decision-making task, we assessed choice accuracy as a function of reward amount across different attentional states (attending to either motion speed or direction). We used another OLS regression, where the dependent variable was choice accuracy (-1 = correct, 1 = correct), and predictors were task state (direction and speed, coded as -1 and 1) and reward value (3 or 5 drops of juice, coded as -1 or 1).

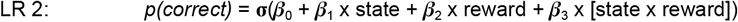

We next analyzed how choice reaction times varied as a function of state and value using general linear regression models. The model (GLM 1) for the state-dependent valuation task regressed reaction times against state (A, B, C, coded as -1, 0, 1) and the absolute difference in values present in the trial.

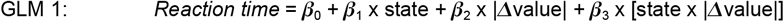

The model (GLM 2) for the perceptual categorization task regressed choice reaction time against task state (direction or speed, coded as -1 or 1) and reward amount (3 or 5 drops of juice, coded as -1 or 1).

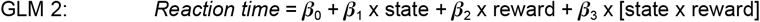

### Neurophysiological recordings

The subject was fitted with a titanium head positioner and imaged in a 3T magnetic resonance imaging scanner. The resulting images were used to generate 3D reconstructions of each subject’s skull and brain areas of interest. We then implanted custom, radiotranslucent recording chambers made of polyether-ether-ketone (PEEK; Gray Matter Research, Bozeman, MT). Each session we acutely lower one 45 mm primate Neuropixels probes (IMEC VZW, Leuven, Belgium) into area 13 of OFC. Electrode trajectories were determined by incorporating the subject’s MRI into computer-assisted drawing software (OnShape, Cambridge, MA) to design and 3D print custom recording grids to lower the probes. We used a Form 4 3D printer (Form Labs, Somerville, MA). In order to mitigate recording drift, the probes were allowed to settle for approximately 60 minutes after insertion, prior to the start of the behavioral task. We configured the probes to record from 384 active channels in a contiguous block at the tip, allowing dense sampling of neuronal activity along a 3.84 mm span. Neuronal activity was filtered and digitized for action potential bands (300 Hz high-pass filter, 30 kHz sampling frequency) and local field potential (1 kHz low-pass filter, 2.5 kHz sampling frequency). Activity was monitored during experimental sessions and saved to disk using Spike GLX (https://billkarsh.github.io/SpikeGLX/).

Spiking in the action-potential band was identified and sorted offline using *Kilosort4* (43). Because each Neuropixels headstage samples the data with a slightly different system clock, it was necessary to map all physiological and task-event time series into a common timeline. We accomplished this by broadcasting a 5 Volt, 1 Hz square wave to a unique reference channel on the Neuropixel probe (channel 385) and the non-neural acquisition board. We detected and stored the times of the rising phase of the 5 V wave on each cycle and then used regression to interpolate all task events into the timeline of the probe. To be included for further analysis, neurons had to be present for >90% of the experimental session, have a mean firing rate over the course of the entire session >1 Hz, and <0.1% of spikes occurring within 1 ms of another isolated neuron’s spikes (that is, <0.1% interspike interval violations).

### Analysis of single neuron tuning

To understand whether the OFC encodes state and value during each task, we performed a sliding-window regression analysis (GLM 3 and GLM 4) assessing how a given neuron’s firing rate varied as a function of task state, chosen value, and the interaction of state and value. We used a one-hot encoding procedure for each of the state regressors, meaning that, for example, β1 x state_A was coded as “1” for all state A trials and “0” for trials of all other states. The value factor was coded as 1-thru-4 for the value task and 3 or 5 for the perceptual task. Each of the interaction terms was defined as the one-hot vector for a given state multiplied by either the chosen value (for the value task) or the expected reward (for the perceptual task). We used this one-hot encoding approach because without it, any state-value interactions in an ANOVA could reflect image-encoding rather than a state-dependent value code.

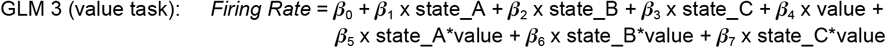

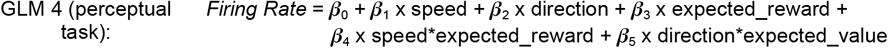

We fit the regression model at each timestep within the trial structure, using 100 ms bins with 25 ms steps between bins. The p-value for each factor for each cell at each timestep was computed and we used an alpha level of *p* <.01. State encoding neurons were defined as those encoding any of the state factors for at least 150 consecutive milliseconds. Value/expected reward neurons were defined as those encoding the value/expected reward factor for at least 150 consecutive milliseconds. State-dependent value neurons were defined as those significantly encoding any of the state*value terms for at least 150 consecutive milliseconds.

### Analysis of task epoch selectivity

To identify neurons that were selectively active during specific task epochs, we performed a one-way ANOVA on firing rates across the three main task epochs (cue, motion stimulus, and choice). For each neuron, we calculated the mean firing rate during a 500 ms window beginning at the onset of each epoch. We first identified neurons which were significantly modulated by task epoch (*p* <.01) and then defined a neuron’s preferred epoch as the one which elicited its highest mean firing rate.

### Population decoding of state-dependent values

To assess the population dynamics within OFC during each task, we trained linear discriminant analysis decoders to classify state-value conjunctions. We used a similar decoding framework for both tasks: for the state-dependent value task, the decoder classified 12 classes (4 reward values x 3 states); for the perceptual decision-making task, the decoder classified 4 classes (2 reward outcomes x 2 cued features). To ensure robust single-trial resolution of our decoders, we used a bootstrapping approach where on each bootstrap 90% of single-saccade trials were randomly selected for training. We then balanced the training data, equating the number of trials where a given value-state combination was selected. The remaining trials were used for testing. This procedure ensured that the composition of our training data did not introduce bias into the decoder. We repeated this balanced train-test split 1000 times at each time step and then averaged over instances where the same trial was present in the testing set across bootstraps. Decoder accuracy was extracted for each time point (25 ms bin) for each trial. We empirically identified the null hypothesis of these decoders (that they detect nothing but noise) by repeating our bootstrapping procedure but shuffling the training labels on each bootstrap.

## Results

To examine OFC’s role in value-based and perceptual decisions, we trained a monkey (Subject K) to perform both a value-based choice task (“value task”) and a perceptual categorization task (“perceptual task”) while conducting Neuropixel recordings in OFC. The recordings were made sequentially with the value task occurring first and then 3 months later the perceptual task. We designed the perceptual task so that its structure paralleled the value task’s (**Fig. 1**).

**Figure 1.**
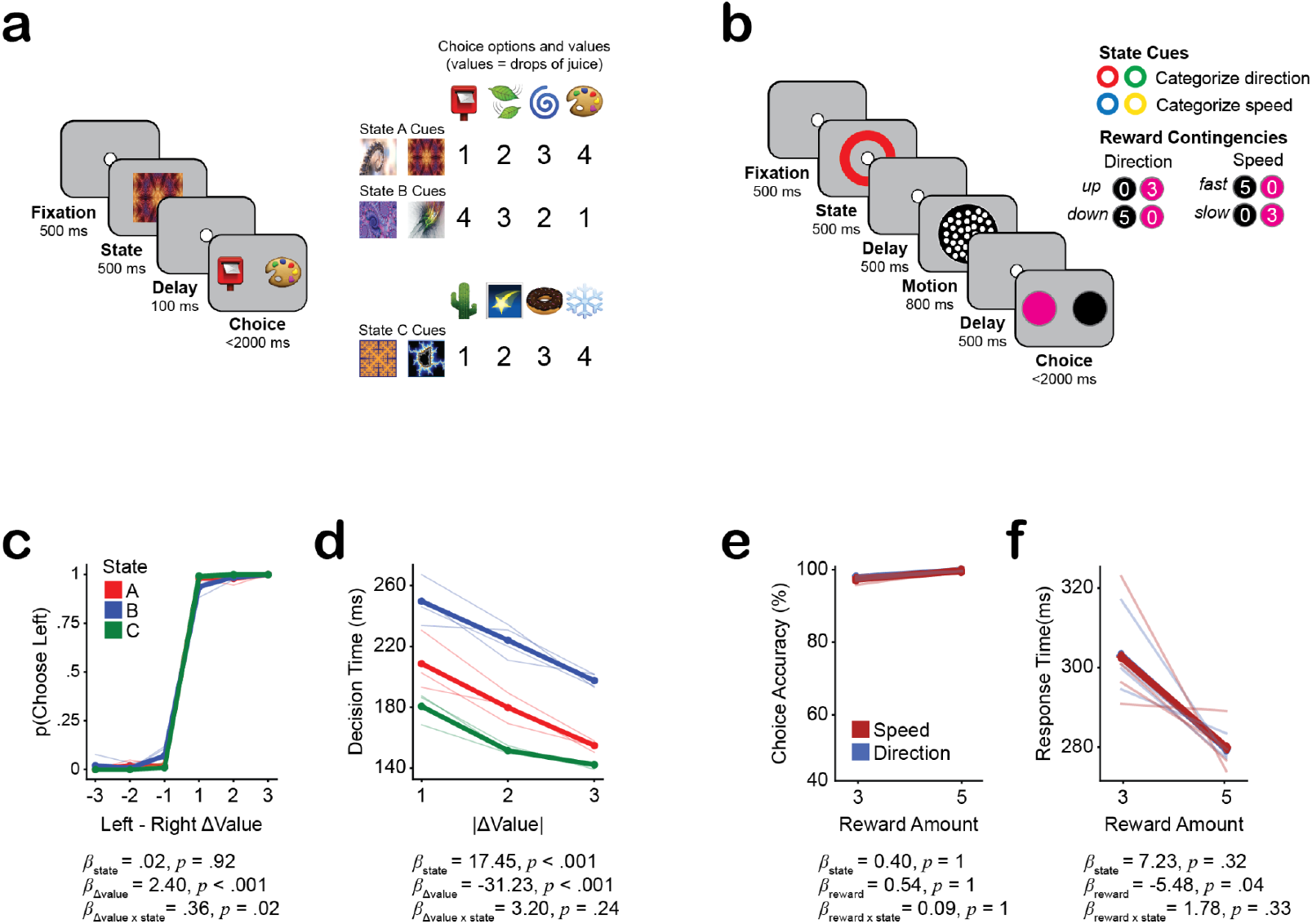
Task designs and behavioral performance. (a) Value based choice task design. The subject initially fixated and was then cued as to which task state the current trial would take place in. Then, after a delay, the subject was presented with either one (forced choices) or two (free choices) options. In states A and B, the rewards associated with the same stimuli changed according to the cued task state; in state C, the rewards associated with the choice options remained constant. We used two cues per task state in order to decorrelate neural responses to the sensory particulars of the stimuli from the meaning of the task state. 75% of trials were free choices, where the subject chose between two options. 25% of trials were forced choices, where only one option was presented and the subject was required to make a saccade to it. (b) Perceptual decision task design. The subject initially fixated and was then cued to attend to either the speed or direction of the upcoming motion stimulus. After a brief delay, a motion stimulus was presented. After a final delay, the subject reported whether the dots were moving up or down (if cued to attend to direction) or whether the dots were moving fast or slow (if cued to attend to speed). Different categories of motion were associated with different reward levels such that correctly categorizing “fast” and “downward” motion paid 5 drops of juice whereas correctly categorizing “slow” and “upward” motion paid only 3 drops of juice. Incorrect categorizations yielded no reward. We used two cues per task state in order to decorrelate neural responses to the sensory particulars of the stimuli from the meaning of the task state. (c) Decision accuracy as a function of option value differences in the value-based choice task. Thin lines are single sessions (n = 4), and thick lines are mean values. The subject’s choice was well predicted by the difference between values. (d) Decision times in the value-based choice task are influenced by expected reward. Faster reaction time occurred in those trials with a greater value difference. (e) Choice performance is not affected by expected reward in the perceptual decision task. Thin lines are single sessions (n = 4), and thick lines are mean values. (f) Decision times in the perceptual task are influenced by expected reward. The subject responded faster when correctly reporting a high-reward category.

In the value task, the subject was trained to choose between different visual stimuli (emojis) to maximize a juice reward associated with different juice amounts. We previously used this task to explore how the OFC encodes state-dependent values (9,16). Critically, the exact amount of juice reward depended on which of 3 task-states was in effect. Each task state was associated with a different reward schema (**Fig. 1a**). Each trial began with a cue indicating which of the three reward schemes (A, B, or C) the upcoming decision should be evaluated within. After a brief delay, the monkey was presented with either one (forced choice, 25% of trials) or two options (free choice, 75% of trials). For states A and B, the same choice stimuli were used, introducing conflict by altering the reward outcome depending on the cued state. In contrast, state C used a distinct set of choice stimuli – this served as a control condition where there were no conflicting value-mappings.

In the perceptual task (**Fig. 1b**), the subject was trained to categorize either the speed or direction of a dot-motion stimulus. Following an initial fixation period, the subject was cued whether to attend to (and thus categorize) the direction or speed of the upcoming motion stimulus. Following a brief delay, the subject was then shown the motion stimulus, during which the monkey had to maintain fixation. The subject was then presented with two decision-targets which allowed the animal to report what he had seen during the motion stimulus epoch. The monkey was trained to report fast motion and downward motion by saccading to a black, speckled stimulus and, conversely, was trained to report slow motion or upward motion by saccading to a pink circle stimulus. Both speed and direction were present in every trial and we over-represented the “conflict” trials where the animal categorized the same motion stimulus differently depending on whether it was currently attending to speed or direction. Each category was associated with a different reward amount such that correctly categorizing either fast motion or downward motion led to a large juice reward (5 drops of apple juice) whereas correctly categorizing either slow motion or upward motion led to a smaller reward (3 drops of apple juice). Incorrect categorization led to no reward. This design enabled us to dissociate the perceptual-basis of the decision (categorizing the motion) from reward associated with the category. This was important in order to test whether OFC encoded the expected reward associated with a given perceptual category independently of the stimulus itself.

The subject displayed a high proficiency in both tasks (**Fig. 1c-d**), selecting the best option 98% of the time during free choices of the value task and correctly categorizing 97% of the time in the perceptual task. In the value task, the monkey selected the higher value option more frequently regardless of spatial position, and exhibited faster response time for trials with larger option value differences. In the perceptual task, choice accuracy was nearly perfect across both attentional dimensions (direction 95%, speed 98%). Logistic regression analysis showed that choice accuracy did not differ across task states (*ß* = 0.4, *p* >.9), expected reward amount (*ß* = 0.54, *p* >.9), or their interaction (*ß* = 0.09, *p* >.9). However, reaction times in the perceptual task were slightly faster for trials where the subject successfully classified a high-reward feature (e.g. downward motion when direction was relevant). This indicates that the subject’s response behavior was influenced by expected reward even though the decision of what to report was not determined by reward.

Both tasks involved flexibly responding to the same stimuli differently and for different reward amounts, depending on the cued task state. Next, we examined single unit responses via Neuropixel recordings in the OFC as the subject performed both tasks. We initially recorded neural activity during the value task and then immediately re-trained the animal to perform the perceptual decision task. Retraining took approximately 3 months. We used the same recording chamber and recording coordinates as well as the same Neuropixel probe for both sets of recordings (**Fig. 2a-b**).

**Figure 2.**
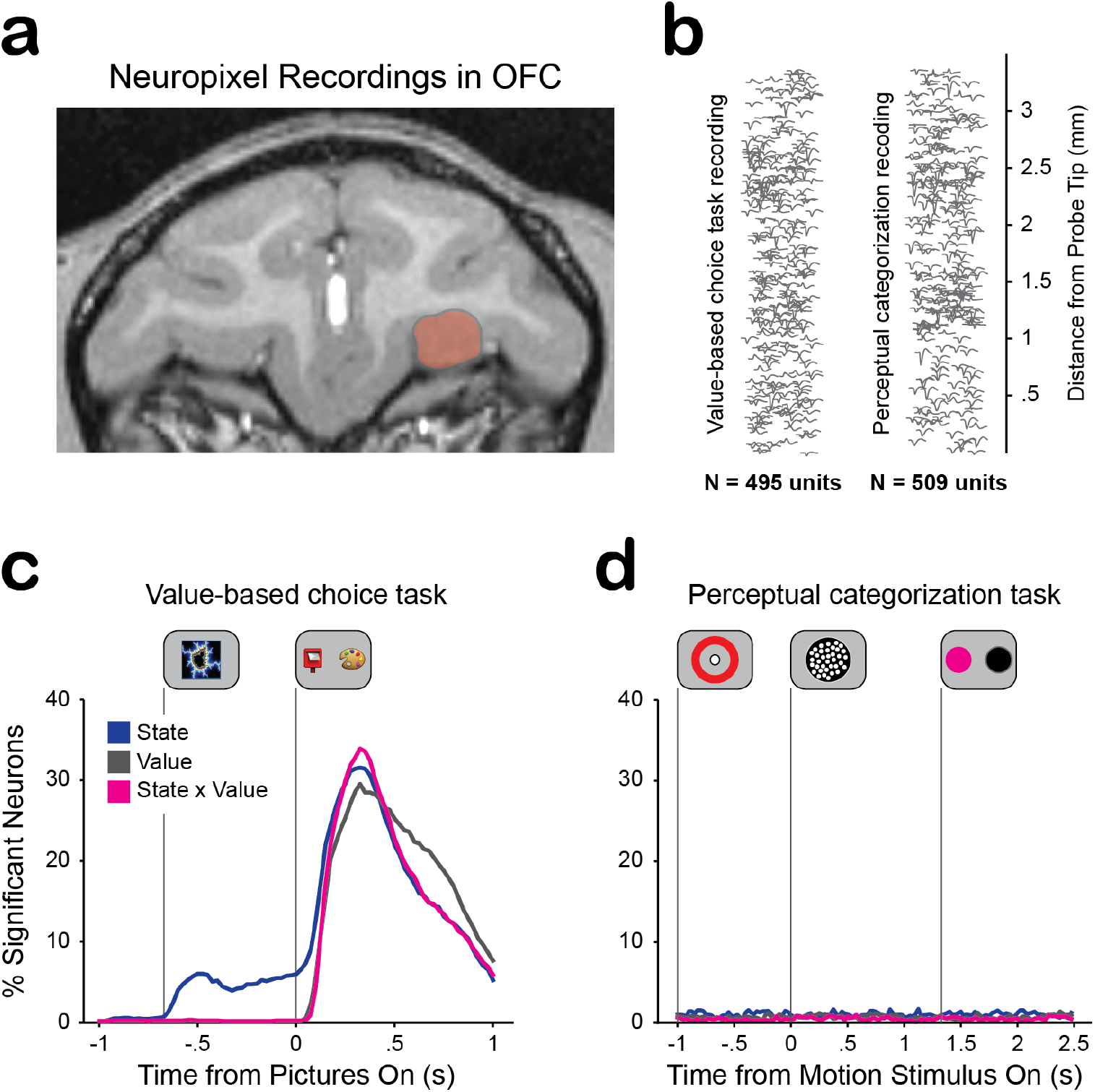
Recording locations and single neuron encoding results. (a) Coronal MRI section showing the recording location in OFC for both tasks. (b) Mean waveforms of hundreds of simultaneously recorded individual OFC neurons localized onto the surface of the Neuropixels probe. These are two representative sessions, one for each task. (c) Percentage of neurons significantly encoding State, Value, or their interaction via sliding window regression. During the state cue epoch, cells began encoding the state while value was found encoded beginning in the choice epoch. (d) Same analysis as **c**. but for the perceptual categorization task. Using binomial tests, we did not observe any significant proportion of OFC neurons encoding any task feature during any epoch of the task.

We report the results of 4 recording sessions per task, where we obtained large numbers of well-isolated single neurons (n = 1528 neurons in the value task; n = 1427 neurons in the perceptual categorization task). To assess encoding of task-relevant variables, we first binned firing rates in 100 ms bins and then used a sliding window regression approach that assessed how firing rates varied as a function of task state, the reward of the chosen option, and the interaction of these factors (see **Methods**). Consistent with our previous report (9), OFC was strongly engaged in the value task with neurons encoding both the task state itself (during the state cue epoch) and all decision-relevant variables (state, value, state x value) of the chosen option during the choice epoch (**Fig. 2c**). In contrast, a minimal number of neurons encoded any factor during the perceptual task (**Fig. 2d**). This indicates that OFC was strongly engaged by the value task but not during the perceptual task.

OFC neurons were not silent during the perceptual task, despite not differentiating decision-relevant features. The majority (1169/1427, 82%) showed significant modulation across task epochs (as assessed with a one-way ANOVA, *p* <.01, see **Methods**). We defined a neuron’s preferred epoch as the one eliciting the highest mean firing rate. For example, **Figure 3a** shows two neurons that were driven by the onset of the motion stimulus but did not differentiate speed, direction, or expected reward. These responses cannot be attributed to purely visual or sensory processing, as the same neurons failed to respond to other stimulus-onset events (e.g., cue onset or choice target onset). Overall, of the 1169 epoch-selective neurons, 557/1169 (47%) preferred the cue epoch, 420/1169 (36%) preferred the motion epoch, and 192/1169 (16%) preferred the choice epoch. As shown in **Figure 3b**, OFC motion-selective units exhibited the strongest responses. This suggests that OFC neurons may be tracking progression through the task structure, consistent with recent state-based, cognitive map theories of OFC function (12,14,17,18). Critically, however, this task-structure selectivity occurred in the absence of any encoding of decision-relevant variables – a stark contrast to the robust representation of state, value, and their interaction observed during the value-based decisions (**Fig. 2c**).

**Figure 3.**
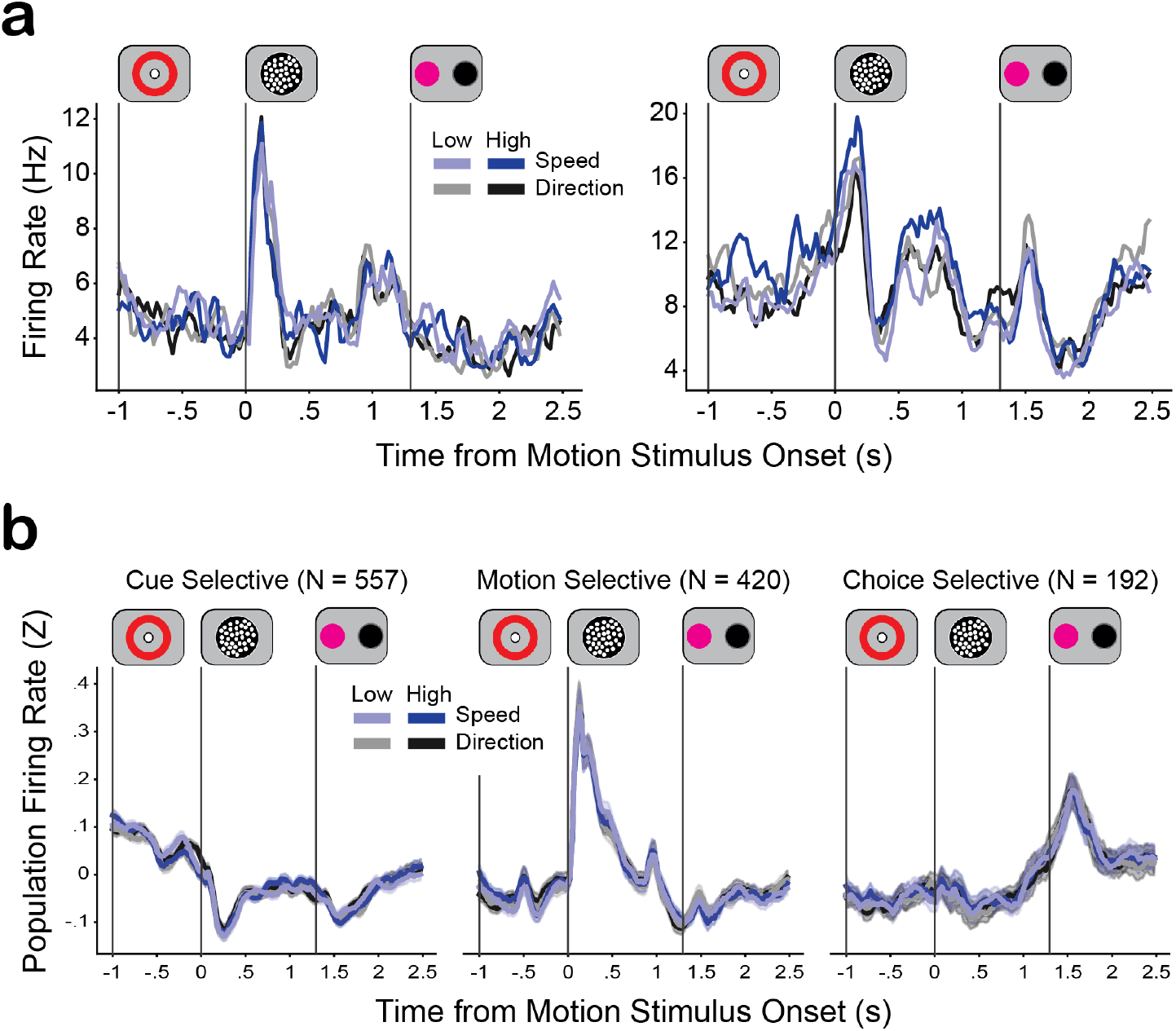
OFC neuron firing rates during the perceptual categorization task. (a) Two OFC neurons that were driven by the onset of the motion stimulus but did not differentiate decision-relevant task features. (b) Trial-averaged firing rates for populations of neurons identified as cue, motion, or choice selective. Neurons identified as significantly modulated (*p* <.01) by task epoch by assessing their firing rates during each task epoch with a one-way ANOVA. A neuron’s preferred epoch was defined as the one which elicited the neuron’s highest firing rate. These traces show the trial-averaged grand mean across neurons identified as preferring each task epoch. Thick lines denote grand means and the shading denotes bootstrapped 95% confidence intervals. We note that the motion-selective population showed the greatest modulation.

Although we could not identify decision-relevant representations at the single-neuron level during the perceptual task, it is possible that such information could be represented at the population level through distributed patterns of activity across neurons. To test this possibility, we examined whether state-dependent values could be decoded from population activity (**Fig. 4**). We used our established procedure of decoding values with single trial resolution (9,19,20). Briefly, we trained linear discriminant analysis classifiers to decode state-value combinations from neural activity. For the value task, a 12-way decoder was trained (4 values x 3 states). For the perceptual task, we trained a 4-way linear discriminant classifier (2 reward levels x 2 attentional states). To obtain robust single-trial estimates of decoded state-dependent values, we used a bootstrapping approach where, within each bootstrap, a random 90% of the trials were used as the training set and the remaining 10% used for testing. We repeated this procedure 1000 times and averaged over instances where the same trial appeared in the testing set multiple times. This yielded a robust measure of state-value coding at each moment during each individual trial. We also generated null distributions by shuffling the training labels and then repeating the bootstrapping procedure.

**Figure 4.**
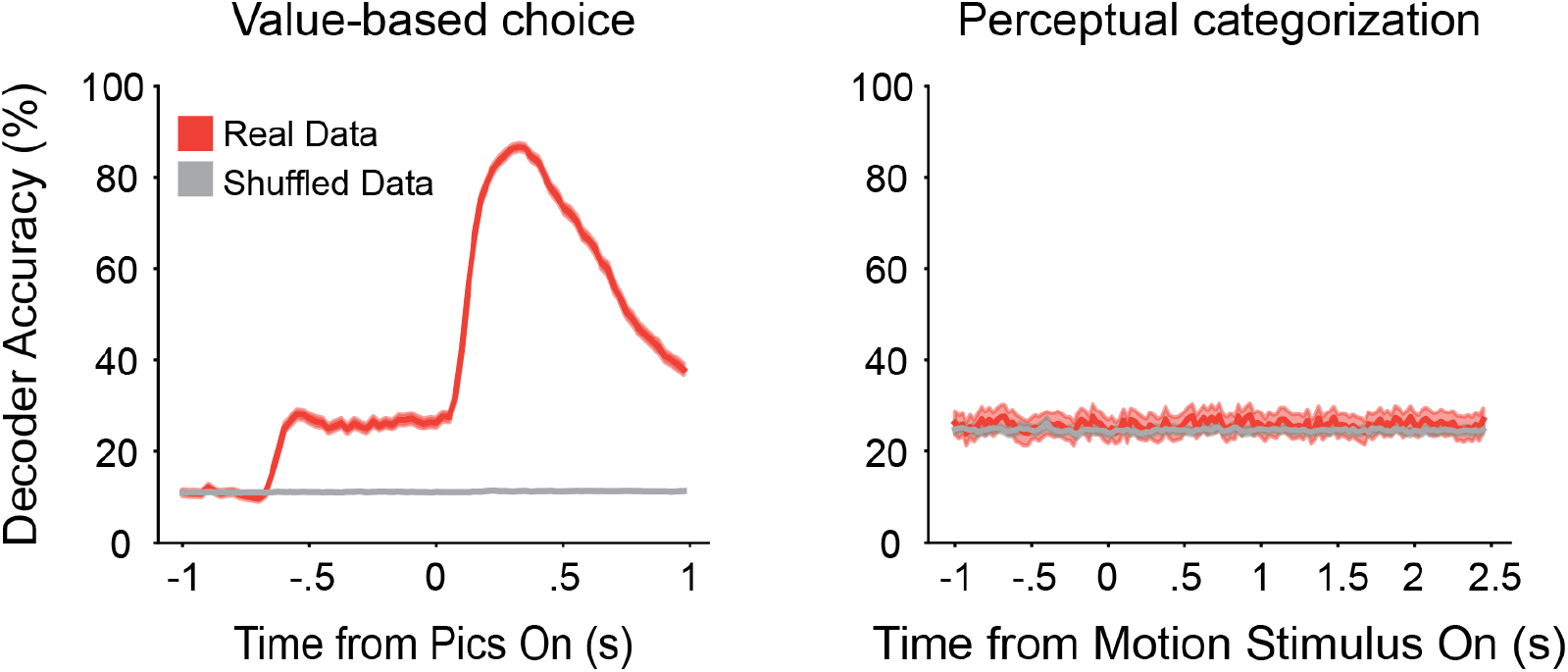
Population decoding of state-dependent value across tasks. a. Average decoder performance over a single trial. Decoder was trained to identify the state-dependent value combination for a single trial based on the neural data. The red line is a decoder trained on unshuffled data while the grey line is a decoder trained on data with shuffled labels. Left is the value-based choice task and right is the perceptual categorization task. For the value-based choice task, decoder performance gradually rose upon presentation of the state cue and then sharply increased upon presentation of the choice options. The perceptual categorization decoder performed comparably to the shuffled data control.

Consistent with the single neuron results, we found robust state-dependent value decoding during the value task. In contrast, the decoder during the perceptual task never differed from the null distribution, indicating that the OFC population did not encode any task-relevant decision variables when the animal made perceptual decisions.

## Discussion

We found that the OFC was robustly activated during a value-based choice task but showed no significant engagement during a perceptual categorization task. Notably, recordings were conducted in the same animal, targeting the same OFC recording sites, using the same Neuropixel probes. We reliably recorded several hundred well-isolated neurons per session during both tasks. This stark difference in OFC’s engagements, may reflect the difference in the cognitive processes underlying value-based choices and perceptual decisions: value-based choices require comparing and selecting between options based on their subjective worth, while perceptual decisions involve categorizing and reporting sensory information. Our finding of selective involvement of OFC in value-based decisions complements an earlier human lesion study (15) and suggests that different neural systems mediate different types of choices, rather than a single unified decision-making network processing all decisions regardless of their underlying cognitive demands.

An alternative possibility is that our results stem from the use of different types of stimuli across the tasks: in the value task, the monkey chose between emojis associated with different rewards; in the perceptual task, the monkey categorized a motion stimulus. These different stimulus types are thought to be mediated by different visual circuits – with object-like emojis being processed in the ventral visual stream (21–24) and motion being processed by the middle temporal area in the dorsal stream (24–28).

OFC receives strong, direct input from the entire ventral stream and only sparse input from the dorsal stream (23,29,30). Thus, one possibility is that OFC’s lack of involvement in the perceptual task could reflect that the task depends on evaluating a sensory input that OFC could only receive through other, intermediate areas.

However, several lines of evidence argue against this interpretation. First, a number of studies have reported OFC engagement in tasks where subjects assigned value on the basis of visual motion and other sensory features associated with the dorsal stream. For example, Moneta et al. (17) found robust OFC value coding when subjects chose between different motion stimuli where different directions were associated with different rewards. Similarly, Bongioanni et al. (31) found clear OFC value coding when subjects made decisions based on the density of dots (where greater density corresponds to greater value). Moreover, a number of studies have documented zones where the dorsal and ventral streams converge (23,32–34), including in the parahippocampal formation (32,35–37), which is another major input to OFC (38,39). These findings indicate that the OFC is capable of generating value predictions based on sensory features associated with the dorsal stream, including visual motion, as used in the present experiment. This suggests that instead of differences in upstream sensory processing and routing, our results are better explained by differences in the underlying cognitive demands of perceptual and value-based decisions.

A key finding from our study was that reaction times in both tasks were influenced by the amount of expected reward. We observed faster response times for decisions that yielded higher rewards, even in the perceptual task where OFC activity was minimal. This suggests that reward-related information is still processed during perceptual decisions, but outside the OFC. Our results complement Gore et al.’s (4) recent results showing that OFC inactivation has no effect on behavior when decisions do not require value-maximization. In their study, one group of rats made value-based choices between two options and another group was conditioned to respond to individually-presented stimuli that predicted different amounts of reward. As in our study, both groups responded faster to higher-reward options. Inactivating the OFC of the rats making value-based choices degraded this reward-response correlation: response times flattened and became decorrelated from expected reward. Strikingly, OFC inactivation had no effect on response times in the Pavlovian group - continuing to respond faster to high-reward predicting stimuli and slower to low-reward predicting stimuli. Similarly, Oyama et al. (40) reported no effect of chemogenetically silencing the OFC of monkeys performing a task where the outcomes were binary (simply rewarded or not) but these same monkeys’ behavior was impaired in a decision-task where the monkeys chose between different options which paid different amounts of reward (juice). Our findings complement this growing body of evidence suggesting that OFC is engaged when subjects must simulate and compare the outcomes associated with different options (12–14), which entails a fundamentally different set of cognitive processes from reporting sensory information, as in visual categorization tasks. Our findings highlight a specific functional role of the OFC in value-based but not perceptual decisions.

This suggests that, rather than a single, unified decision network, the brain may recruit specialized sub-systems to support different types of decisions. An important direction for future research is to map the boundaries between specialized and shared decision systems across decision types. If certain brain areas are recruited across multiple types of decisions, it raises the intriguing possibility that a “central executive” mechanism (41,42) may dynamically recruit specialized sub-circuits based on underlying cognitive demands of the decision at hand.

